# The plastid genome characteristics of a moth orchid (*Phalaenopsis wilsoniii*, Orchidaceae)

**DOI:** 10.1101/2021.03.30.437522

**Authors:** KeKe Xia, Ding-Kun Liu, Jie-Yu Wang

## Abstract

Phalaenopsis wilsonii is a typical deciduous species in the horticulturally well-known genus, *Phalaenopsis*. Tshi species is belonging to the section *Aphyllae* in moth orchid, and is endemic to South China. Although the *Aphyllae* section display the deciduous feature that is unique in this genus, their genetic information is still insufficient and limited them as breeding parent in moth orchid. Here, we reported and characterized the complete chloroplast genome for *Phalaenopsis wilsonii*. We found the total size of the chloroplast genome was 145,373 bp, constituting of a large single copy (LSC) region (84,996 bp), a small single-copy region (10,668 bp) and two inverted repeats (IRs) regions (24,855 bp). Based on homologous searching on database, we annotated 76 protein-coding genes, 38 tRNA, and 8 rRNA. The phylogenetic reconstruction revealed that *P. wilsonii* show the closest relationship with *P. lowii* within subgenus *Parishianae*.

## Introduction

Moth orchid (*Phalaenopsis* Blume) is widely used in gardening around the world, and also occupied a larger proportion of orchid industry (Van Huylenbroeck 2018). Recently, the taxonomy of *Phalaenopsis* reached a congruence in perspectives of morphological and molecular evidence, resulting in four subgenera, subgen. *Phalaenopsis, Parishianae, Hygrochilus* and *Ornithochilus* (Kocyan and Schuiteman 2014; Li et al., 2014, 2016). Subgen. *Parishianae* consists of more than a half of the species richness in *Phalaenopsis*, and is also an important breeding resource, however, the genetic data of this section is quite insufficient. So far, only one complete chloroplast genome of species belonging to this subgenus can be found (Wang et al., 2019), and no transcriptome or nuclear genome data have been published. *P. wilsonii* is endemic in China, and is also a typical deciduous *Phalaenopsis* that is belonging to subgen. *Parishianae*. Here, we provide a complete chloroplast genome of *P. wilsonii*, which will facilitate our understanding on *Phalaenopsis* breeding and future designment of molecular markers.

## Methods and materials

The sampling individual of *Phalaenopsis wilsonii* is cultivated in National Orchid Conservation Centre in Guangdong province of China (114°19’01’’E, 22°60’34’’N), and a voucher specimen (noccphal031n) was also deposited in the Herbarium of National Orchid Conservation Centre, Shenzhen, China. A young leaf from the voucher specimen was sampled to extract the total DNA, and sequencing was performed by Illumina HiSeq 2000 platform (Illumina, San Diego, CA). To obtain the chloroplast source reads, we reconstruct the reference database based on all published *Phalaenopsis* plastid genomes, and mapped our clean reads against them to obtain the chloroplast reads for *P. wilsonii*. We used PLATANUS (Kajitani et al., 2014) to construct the initial contigs, and linked the contigs based on the same reads for the scaffolds, resulting in the final complete genome after artificial modification. The BLAST was used to map the reads onto the genome again to confirm the IR boundaries, and the annotation was performed using Geneious 2019.0.3. The resulting complete chloroplast genome of *P. wilsonii* was submitted to GenBank under the accession number of MW218959. The syntenic block detecting and maximum-likelihood phylogenetic reconstruction are conducted by Homblock (Bi et al., 2018) and RAxML, respectively. The matrix contained the whole plastid genomes of *P. wilsonii*, four moth orchids, and also the other ten orchids. *Cattleya crispate* was applied as outgroup according to the topology from Givnish et al. (2015).

## Result and discussion

The total length of *P. wilsonii* chloroplast genome is 145,373bp, which is slightly less than other published moth orchids (from 146,834 bp in *P. lowii* to 148,964 bp in *P. aphrodite* subsp. *formosana*) with the GC content as 36.9%. As in other orchids, the chloroplast genome of *P. wilsonii* consists of a large single copy (LSC) region (84,995 bp) and a small single-copy region (10,668 bp), which were separated by two inverted repeat (IRs) regions (24,855 bp). Overall, 122 genes (contain repeat region gene) were annotated, including 76 protein-coding genes, 8 rRNAs, and 38 tRNAs. As similar to the description in (Chang et al., 2005), all *ndh* genes of *P. wilsonii* were also non-functional, and the *ndhE* was also lost.

Our phylogenetic tree showed a similar topology to Givnish et al. (2015), where the Vandeae presented a sister relationship to Cymbidieae. And *P. wilsonii* was sister to *P. lowii*, and these two species belonging to subgen. *Parishianae* formed the single clade (Figure 1). This complete chloroplast genome of *P. wilsonii* will be the useful information for future phylogenetic studies and conservation in *Phalaenopsis*.

**Figure 1.**
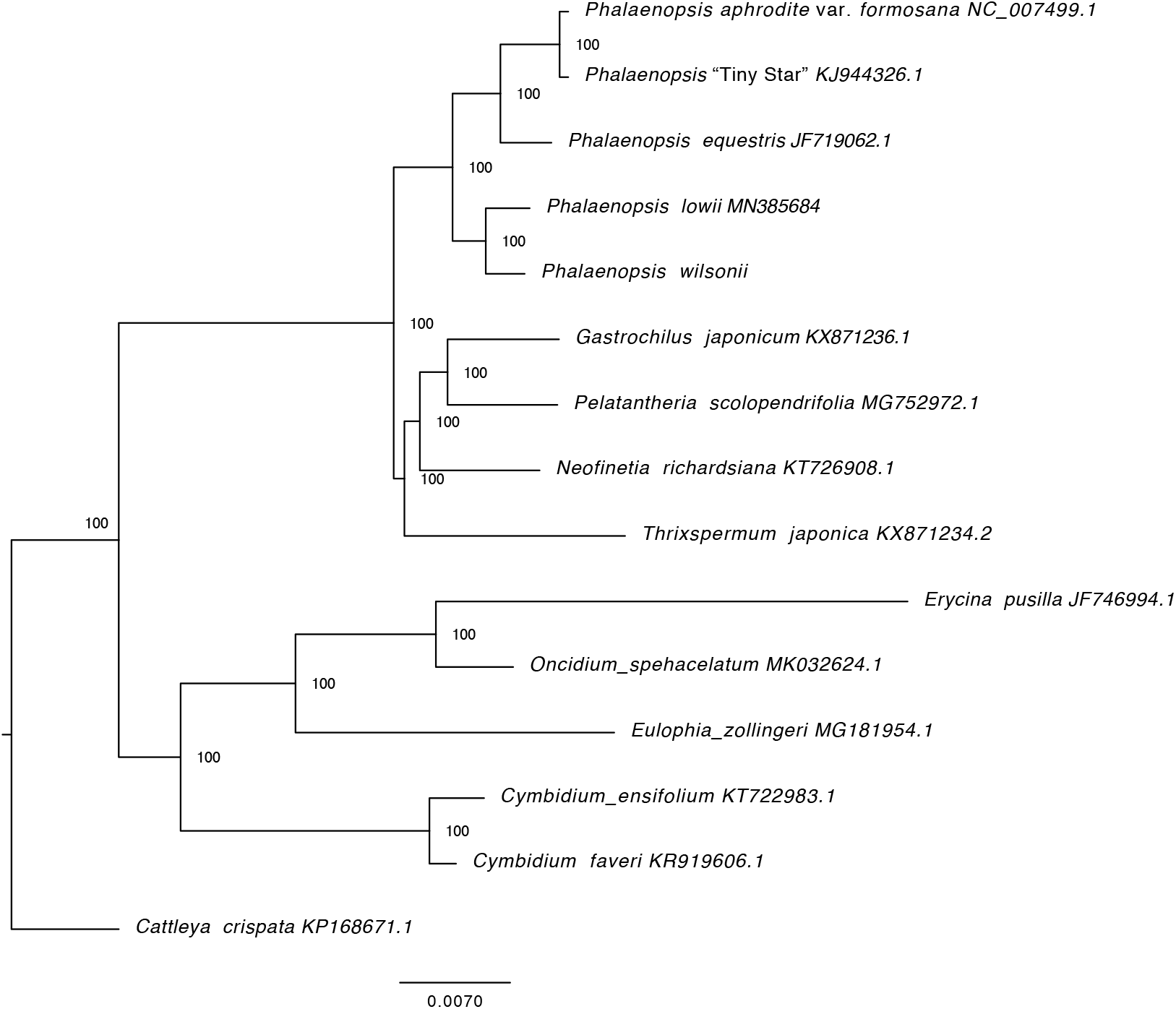
Maximum-likelihood tree reconstructed by RAxML based on complete chloroplast genome sequences from *P. wilsonii*, four *Phalaenopsis* species, and ten other orchids. *Cattleya crispate* was set as outgroup. Numbers on branches are bootstrap support values.

## Acknowledgement

This work was supported by the Science, Technology and Innovation Commission of Shenzhen Municipality under Grant No. JCYJ20170817151501595.

## Authors’ contributions

J.Y.W. and K.K.X. conceived the study; J.Y.W. and K.K.X. obtained the molecular data; J.Y.W. and D.K.L. conducted the data analysis; K.K.X. drafted the manuscript; J.Y.W. and D.K.L. revised the manuscript. All authors provided comments and final approval.

## Data Availability Statement

The data that support the findings of this study are openly available in GenBank of NCBI at https://www.ncbi.nlm.nih.gov, reference number MW218959.

